# Intrinsic maturation of sleep output neurons regulates sleep ontogeny in *Drosophila*

**DOI:** 10.1101/2021.10.14.464413

**Authors:** Naihua N. Gong, An H. Dang, Benjamin Mainwaring, Emily Shields, Karl Schmeckpeper, Roberto Bonasio, Matthew S. Kayser

**Affiliations:** Department of Psychiatry, Perelman School of Medicine at the University of Pennsylvania, Philadelphia, Pennsylvania 19104; Department of Urology and Institute of Neuropathology, Medical Center-University of Freiburg, 79106 Freiburg, Germany; Epigenetics Institute and Department of Cell and Developmental Biology, Perelman School of Medicine at the University of Pennsylvania, Philadelphia, Pennsylvania 19104; Department of Computer Science, University of Pennsylvania, Philadelphia, Pennsylvania 19104; Department of Neuroscience, Perelman School of Medicine at the University of Pennsylvania, Philadelphia, Pennsylvania 19104; Chronobiology and Sleep Institute, Perelman School of Medicine at the University of Pennsylvania, Philadelphia, Pennsylvania 19104

## Abstract

The maturation of sleep behavior across a lifespan (sleep ontogeny) is an evolutionarily conserved phenomenon. Mammalian studies have shown that in addition to increased sleep duration, early life sleep exhibits stark differences compared to mature sleep with regard to the amount of time spent in certain sleep states. How intrinsic maturation of sleep output circuits contributes to sleep ontogeny is poorly understood. The fruit fly *Drosophila melanogaster* exhibits multifaceted changes to sleep from juvenile to mature adulthood. Here, we use a non-invasive probabilistic approach to investigate changes in sleep architecture in juvenile and mature flies. Increased sleep in juvenile flies is driven primarily by a decreased probability of transitioning to wake, and characterized by more time in deeper sleep states. Functional manipulations of sleep-promoting neurons in the dFB suggest these neurons differentially regulate sleep in juvenile and mature flies. Transcriptomic analysis of dFB neurons at different ages and a subsequent RNAi screen implicate genes involved in distinct molecular processes in sleep control of juvenile and mature flies. These results reveal that dynamic transcriptional states of sleep output neurons contribute to changes in sleep across the lifespan.

## Introduction

Across species, sleep duration peaks in early life and declines with age (Jouvet-Mounier et al., 1969; Kayser and Biron, 2016; Roffwarg et al., 1966). Early life sleep is also characterized by differences in sleep architecture compared to maturity. For example, in humans, sleep duration as well as percentage of time spent in rapid eye movement (REM) sleep is highest in newborn infants and decreases with age (Roffwarg et al., 1966). Several lines of evidence point towards the importance of early life sleep for normal neurodevelopment (Blumberg, 2015; Cao et al., 2020; Frank et al., 2001; Jones et al., 2019; Kayser et al., 2014; Marks et al., 1995; Seugnet et al., 2011). Juvenile sleep may thus have characteristics that fulfill specific needs for nervous system development. However, mechanisms underlying sleep ontogeny -- the change in sleep features across development – are largely unknown.

At a fundamental level, the probability of transitioning between sleep and wake influence sleep duration. These transitions are controlled by an interplay between sleep regulatory neural substrates (Artiushin and Sehgal, 2017; Eban-Rothschild et al., 2018; Scammell et al., 2017). In addition, both mammals and invertebrates such as *Drosophila melanogaster* exhibit transitions between distinct sleep stages, which are defined by electrophysiologic and behavioral measurements (Blake and Gerard, 1937; Clancy et al., 1978; Lendner et al., 2020; Nitz et al., 2002; Tainton-Heap et al., 2021; Weber, 2017; Yamabe et al., 2019; Yap et al., 2017). In *Drosophila*, conditional probabilities of activity/inactivity state transitions as well as hidden Markov modeling of sleep/wake substates have proven to be useful, non-invasive methods for probing the neurobiology underlying sleep architecture (Wiggin et al., 2020). Using such approaches towards a detailed analysis of sleep/wake transitions and sleep states in juvenile flies has yet to be explored.

How does the development of sleep-regulatory circuits influence changes to sleep architecture across the lifespan? In flies, maturation of a key sleep circuit in the central complex of the brain contributes to sleep ontogenetic changes. Specifically, juvenile flies exhibit increased activity in sleep-promoting neurons of the dorsal fan-shaped body (dFB) compared to mature flies (Kayser et al., 2014). One factor governing this change in sleep output is the maturation of dopaminergic (DA) inputs that inhibit dFB activity (Liu et al., 2012; Pimentel et al., 2016; Ueno et al., 2012). These DA inputs are both less numerous and less active in juvenile flies, leading to increased dFB activity compared to mature flies (Chakravarti Dilley et al., 2020; Kayser et al., 2014). However, whether sleep-promoting dFB neurons themselves also undergo intrinsic maturation is unknown.

Using a conditional probabilities approach applied to locomotor measurements and hidden Markov modeling of sleep/wake substates (Wiggin et al., 2020), we addressed the question of how sleep architecture differs between juvenile and mature *Drosophila*. We find excess sleep in juvenile flies is driven primarily by a decreased probability of flies transitioning out of sleep. Juvenile flies additionally spend proportionally more time in a deep sleep state compared to mature flies. Activation in mature flies of sleep-promoting neurons defined by *R23E10-GAL4* increases sleep duration, but yields sleep architecture distinct from the juvenile sleep state. Conversely, inhibition of the same dFB neurons in juvenile flies does not result in mature fly sleep architecture. Finally, we find the dFB exhibits distinct molecular signatures across the period of sleep maturation, supporting the idea of an evolving role for the dFB across development. Our results suggest that intrinsic maturation of sleep-output neurons contributes to sleep ontogenetic changes.

## Results

### Juvenile flies exhibit increased deep sleep compared to mature flies

To investigate how sleep/wake transition probabilities differ between juvenile (1 day post-eclosion) and mature (5-7 days post-eclosion) adult flies, we recorded sleep in *iso31* female flies using a high-resolution multibeam Drosophila Activity Monitoring (DAM) system. Consistent with previous studies (Dilley et al., 2018; Kayser et al., 2014; Shaw et al., 2000), we observed greater sleep duration both during the day (ZT0-12) and night (ZT12-24) in juvenile flies compared to mature flies (**Fig 1A**). P(wake) is defined as the probability of transitioning from an inactive to an active state, while P(doze) is the probability of transitioning from an active to inactive state (Wiggin et al., 2020). P(wake) was significantly decreased during the day and the night in juvenile flies (**Fig 1B**), suggesting that increased sleep duration in juvenile flies is driven primarily by a lower probability of transitioning from sleep to wake. P(doze) was also decreased in juvenile flies across the day and night (**Fig 1C**). Previous work has established that P(doze) is less closely correlated with sleep duration than P(wake) (Wiggin et al., 2020), consistent with our observation that P(wake) is decreased in juvenile flies and drives increased sleep duration. We noted more variance in P(doze) in juvenile flies compared to mature when measured during 30-minute windows (**Fig S1**), likely because young flies spend so much time asleep that transitioning from wake to sleep is a relatively rare event over this short period of time. A larger 12-hour window (**Fig 1C, right**) more accurately assesses P(doze), especially in juvenile flies.

**Fig 1:**
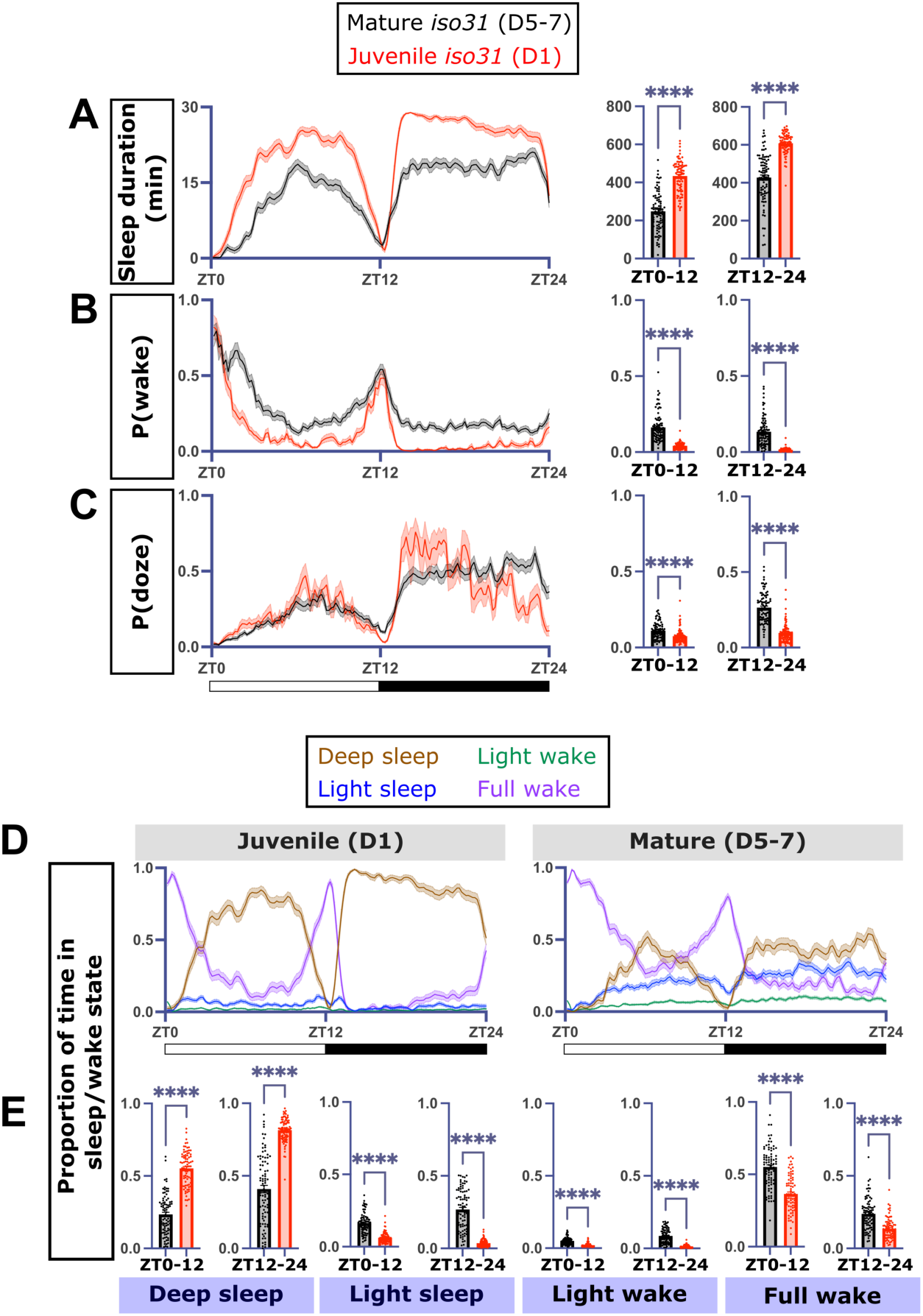
Excess sleep in juvenile flies is characterized by increased deep sleep driven by a decreased probability of transitioning from sleep to wake. A) Sleep duration, B) P(wake), and C) P(doze) in mature (black, n = 87) vs juvenile (red, n = 82) *iso31* flies. Left: sleep metric traces. Right: Quantification of sleep metrics across the lights-on (ZT0-12) or lights-off (ZT12-24) periods. D) Deep sleep (brown), light sleep (blue), light wake (green), and full wake (purple) traces in juvenile (left) and mature (right) *iso31* flies. E) Quantification of proportion of time spent in each sleep stage across the lights-on or lights-off periods (two-tailed T-tests for A-E). For this and all subsequent figures, sleep metric traces are generated from a rolling 30-minute window sampled every 10 minutes unless otherwise specified. For graphs in this figure and all other graphs unless otherwise specified, data are presented as mean ± SEM. \**P* < 0.01, \*\**P* < 0.01, *\*\*\*P* < 0.001, *\*\*\*\*P* < 0.0001.

Next, we asked how sleep/wake stages differ between juvenile and mature flies. In the presence of an arousing stimulus during sleep, juvenile flies are less likely to wake compared to their mature counterparts (Kayser et al., 2014). In *Drosophila*, an increased arousal threshold is indicative of a deeper sleep state (Wiggin et al., 2020), but the proportion of time spent in specific sleep states across the lifespan is unknown. Locomotor recording followed by hidden Markov modeling has been successfully used as a non-invasive method to establish physiologically-relevant sleep/wake substates from DAM system activity measurements (Wiggin et al., 2020). We trained two hidden Markov models (HMMs) with four hidden substates (deep sleep, light sleep, light wake, and full wake) using activity measurements from mature or juvenile *iso31* flies (**Table S1**). To determine whether transition and emission probabilities of the HMMs trained on mature and juvenile datasets (HMM-old and HMM-young) differed, we calculated the probability that HMM-old or HMM-young exactly fit observed activity patterns of each fly. For both juvenile and mature fly datasets, HMM-old and HMM-young each yielded significantly different probabilities (**Fig S2A-B**), suggesting the characteristics of defined sleep/wake substates are dynamic across the lifespan. Applying HMM-old and HMM-young to the datasets yielded minor differences in the proportion of time spent in each of the four substates for both mature and juvenile flies. Despite these distinctions, the trends in substate differences between mature and juvenile flies were the same regardless of the model used (**Fig S2C-F**), showing either model can be generally applied to observe biologically-relevant differences in sleep states between juvenile and mature flies.

We applied the HMM trained on mature fly activity to determine the proportion of time juvenile and mature flies spent in each of the four substates (**Fig 1D)**. Compared to mature flies, juvenile flies spent significantly more time in deep sleep across both the day and night. This proportional increase came at the expense of light sleep, light wake, and full wake (**Fig 1E**). Thus, the propensity for juvenile flies to spend more time in a less-arousable deep sleep state may explain the lower probability of transitioning from sleep to wake.

**Table S1:**
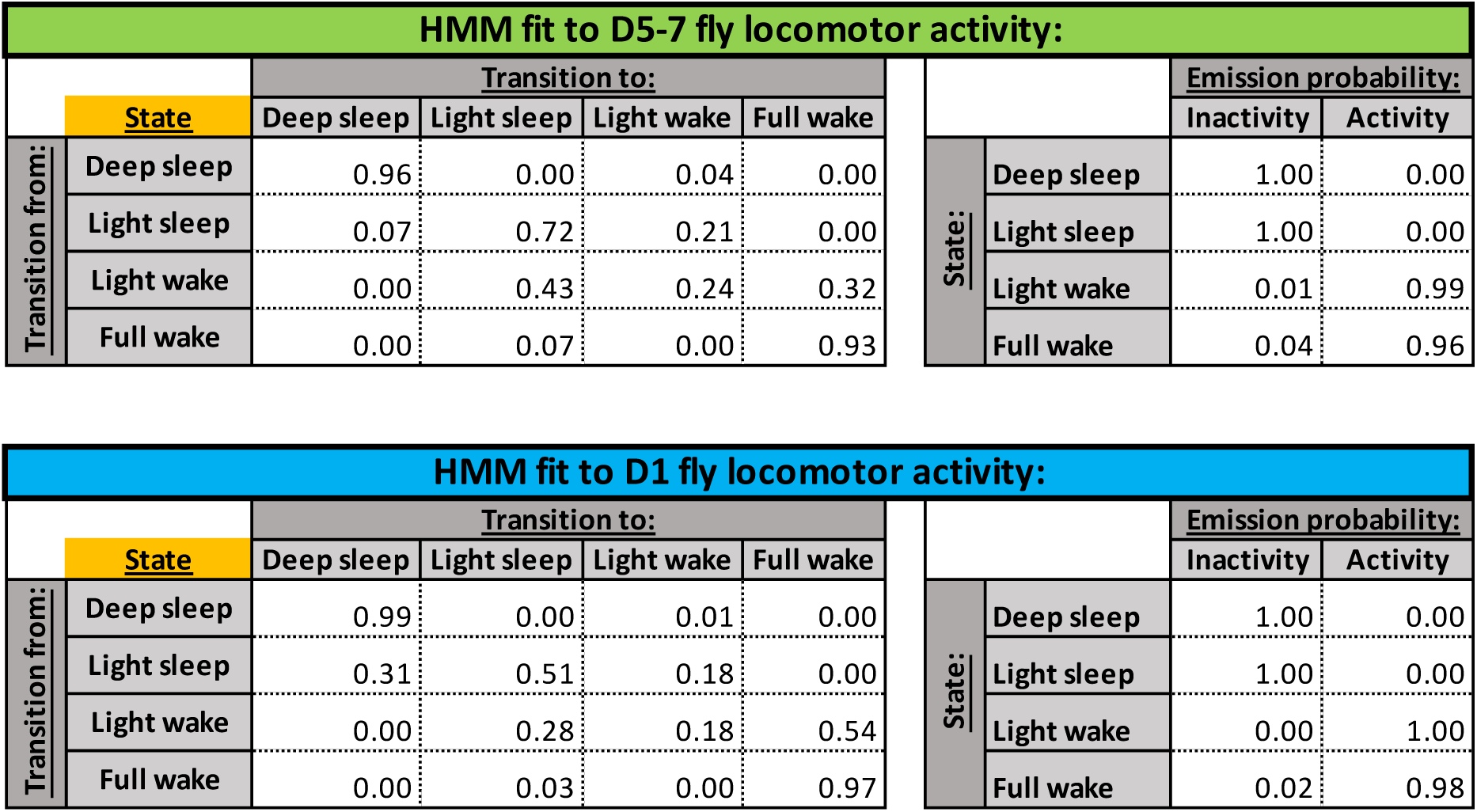
Transition probabilities between hidden states and emission probabilities from each hidden state to observed states for HMM trained on mature (top) and juvenile (bottom) *iso31* fly locomotor data collected using the DAM5H multibeam system. HMMs were trained on transitions from each fly (n = 87 for mature flies, n = 82 for juvenile flies; total 1439 transitions per fly per 24 hours).

**Fig S1:**
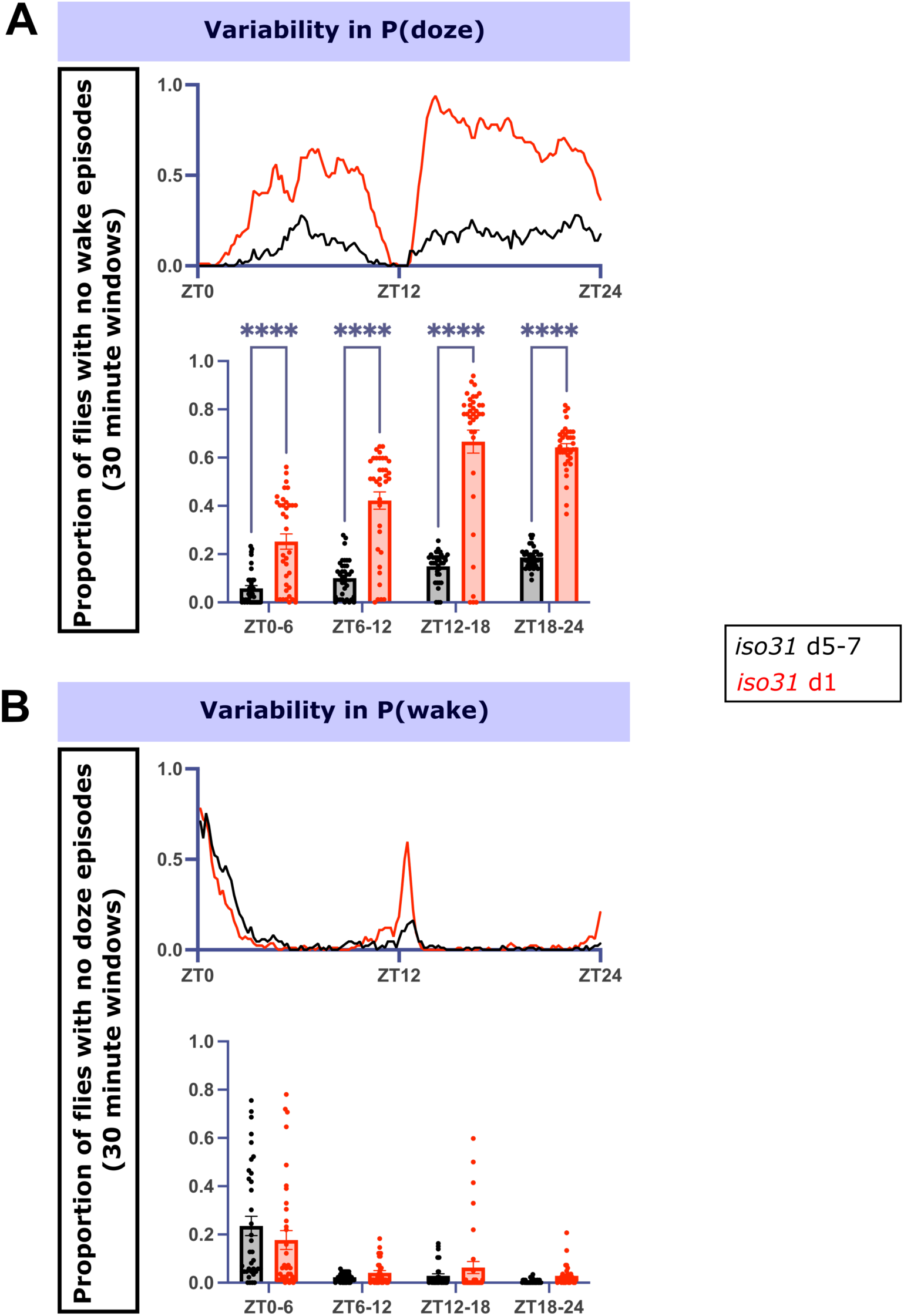
Increased variance in 30-minute windows of P(doze) in juvenile flies compared to mature flies. Proportion of undefined A) P(doze) and B) P(wake) values across 24 hours in mature (black, n = 87) and juvenile (red, n = 82) flies shown in Figure 1. Top traces are a rolling 30-minute window sampled every 10 minutes, Bottom graphs show the average proportion of undefined values per 30-minute window across 6-hour intervals.

**Fig S2:**
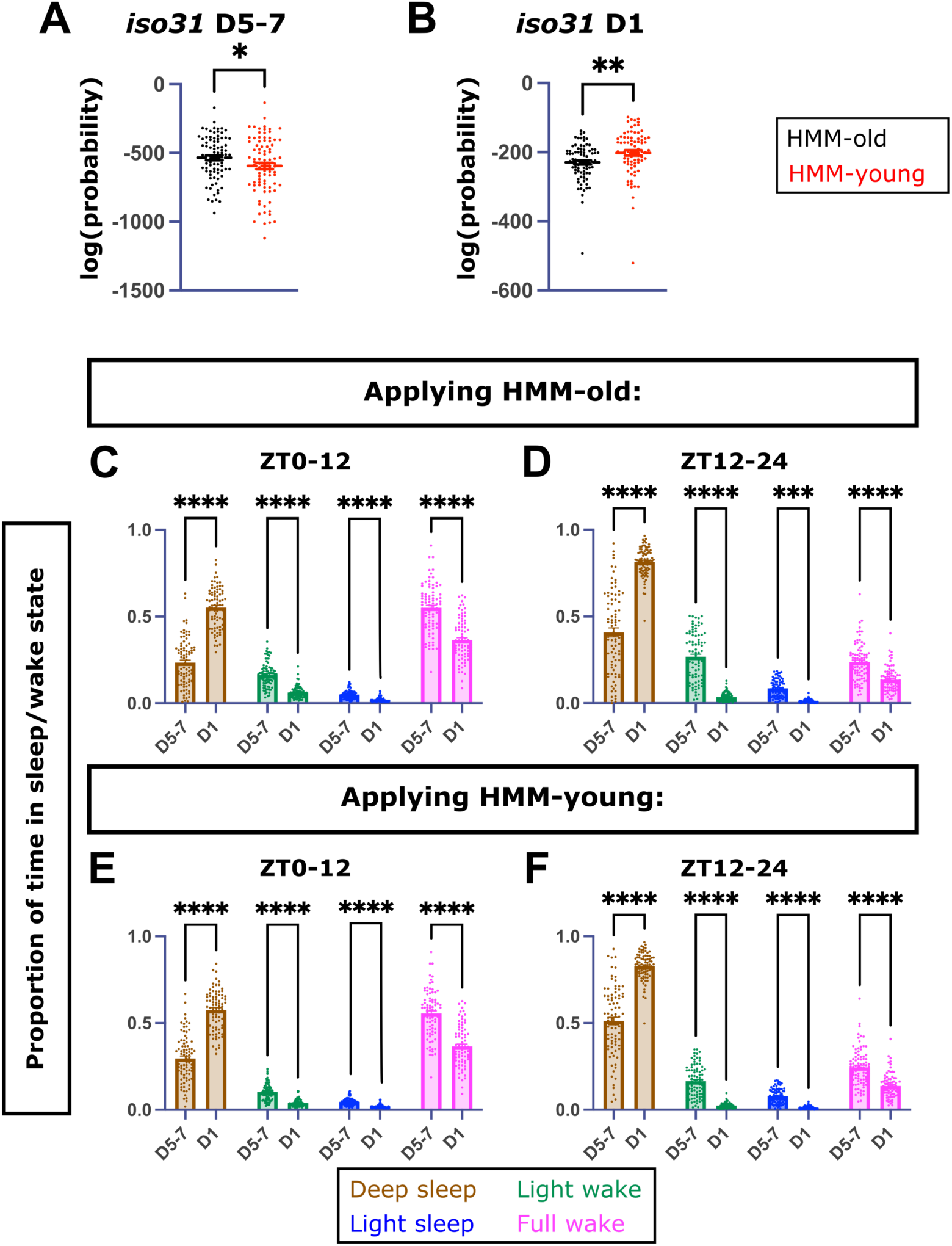
HMMs trained on mature vs juvenile *iso31* fly locomotor datasets have significantly different characteristics. Log(probability) of observing a given sequence of locomotor behavior in A) mature or B) juvenile *iso31* flies by applying HMM-old (black, n = 87) or HMM-young (red, n = 82) (two-tailed T-tests). Proportion of time spent in each sleep/wake hidden state in mature vs juvenile flies from ZT0-12 and ZT12-24 when applying C, D) HMM-old or E, F) HMM-young (two-way ANOVA with post-hoc Sidak’s multiple comparison test).

### The juvenile sleep state is distinct from rebound sleep in deprived mature flies

Our data show that juvenile flies exhibit decreased P(doze) (**Fig 1C**). This change is distinct from previous studies of rebound sleep in mature flies after deprivation, which is characterized by increased P(doze) (Wiggin et al., 2020). To test this distinction directly, we mechanically sleep deprived mature *iso31* flies from ZT12-24, and recorded rebound sleep during ZT0-12 (**Fig 2A**). Deprived mature flies slept significantly more than control mature flies and juvenile flies until ZT6-12, when sleep duration tapered off to non-deprived mature fly levels (**Fig 2B**). P(wake) in rebounding mature flies was significantly decreased compared to control mature flies from ZT0-6 (**Fig 2C**), while P(doze) was increased during ZT0-9 compared to mature controls. Of note, even though sleep duration in deprived mature flies and juvenile flies was comparable around ZT3-9 (**Fig 2B**), P(doze) in deprived mature flies remained elevated across the entire ZT0-12 period compared to juvenile flies (**Fig 2D**). Finally, we assessed sleep substates (**Fig 2E-H**) and found deep sleep was significantly increased in rebounding mature flies, although the deep sleep changes did not persist across the entire day as in juvenile flies (**Fig 2E**). These results support the idea that juvenile fly sleep is a unique state that is different from mature fly homeostatic sleep rebound.

**Fig 2:**
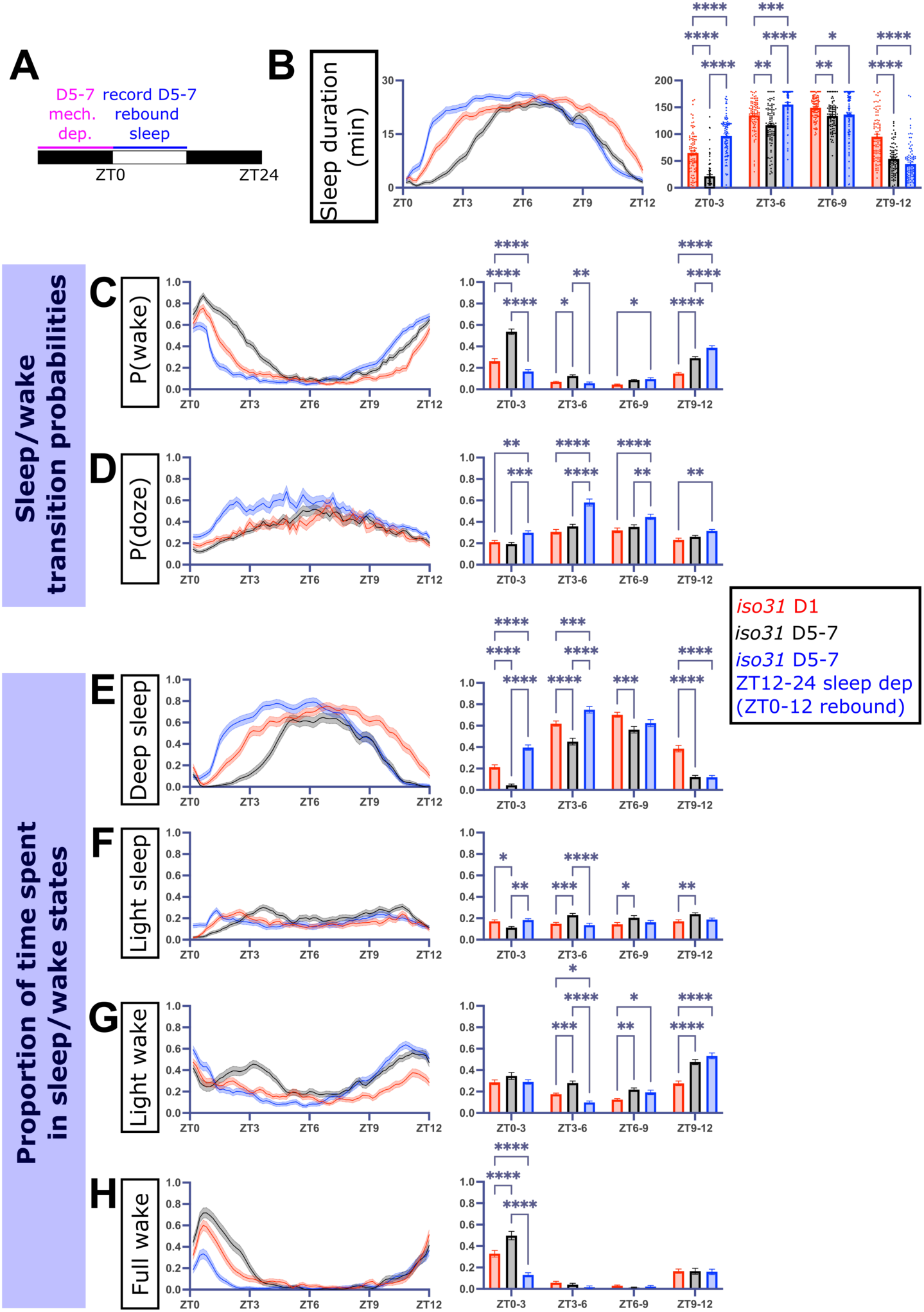
The juvenile sleep state is distinct from homeostatic sleep rebound in mature flies. A) Schematic of deprivation period and period of recorded rebound sleep in mature flies. B) Sleep duration, C) P(wake), D) P(doze), and proportion of time spent in E) deep sleep, F) light sleep, G) light wake, and H) full wake in rebounding mature *iso31* flies (blue, n = 90) compared to non-deprived *iso31* mature flies (black, n = 85) and juvenile *iso31* flies (red, n = 90) (one-way ANOVA with post-hoc Tukey’s multiple comparison test). For figures B-H, data shown is from ZT0-12 after overnight ZT12-24 deprivation. Left: sleep metric traces. Right: quantification of sleep metrics binned into 3-hour windows across ZT0-12.

### Sleep-promoting dorsal fan-shaped body neurons exhibit differential function between juvenile and mature flies

During sleep rebound following deprivation, the dFB exhibits increased activity in mature flies (Donlea et al., 2014). Since the dFB is more active in juvenile flies (Kayser et al., 2014), we next asked whether activation of the dFB in mature flies results in a juvenile-like sleep state. We thermogenetically activated a sleep-promoting subset of dFB neurons using *R23E10-GAL4* (Donlea et al., 2018, 2014) to drive a heat-sensitive cation channel, *UAS-TrpA1* (Hamada et al., 2008) (*R23E10-GAL4>UAS-TrpA1*) in mature flies. Compared to a baseline 24 hours at 22°C, raising the temperature to 31°C significantly increased sleep during the day and the night compared to genetic controls in mature flies (**Fig 3A, E**). Activation of dFB neurons decreased P(wake) during both the day and the night, without changes to P(doze) during the day and increased P(doze) at night compared to one genetic control (**Fig 3B-C, F-G**). Finally, activation of dFB neurons also increased deep sleep in mature flies while decreasing the amount spent in the three other sleep/wake substates (**Fig 3H-K)**. Thus, dFB neuron activation increases sleep in mature flies and mirrors some aspects of juvenile sleep; however, this manipulation also increases P(doze), distinct from the decrease in P(doze) normally observed in juvenile flies.

**Fig 3:**
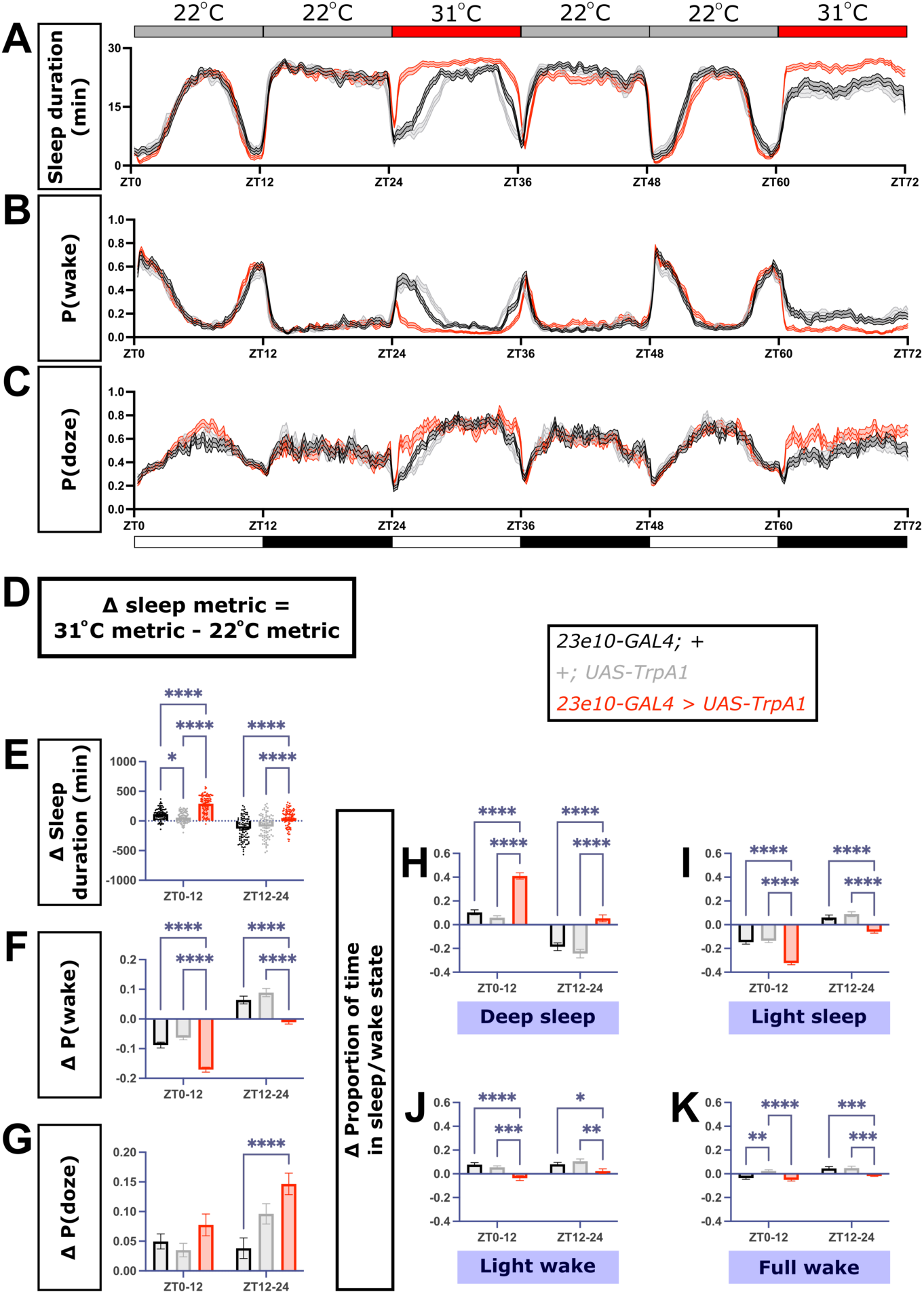
Thermogenetic activation of *R23E10-GAL4+* neurons in mature flies does not fully recapitulate the juvenile sleep state. A) Sleep duration, B) P(wake), and C) P(doze) traces in *R23E10-GAL4>UAS-TrpA1* (red, n = 102) flies and genetic controls (black, n = 97 and gray, n = 85). Gray bars at the top denote periods at 22°C, while red bars denote periods at 31°C. D) Formula used to calculate changes in sleep metrics. To account for differences in baseline sleep metrics between different genotypes at 22°C, changes in sleep metrics for individual flies was calculated. Change in E) sleep duration, F) P(wake), and G) P(doze) across ZT0-12 and ZT12-24. Changes in the proportion of time spent in H) deep sleep, I) light sleep, J) light wake, and K) full wake in the setting of thermogenetic *R23E10-GAL4* neuron activation (one-way ANOVA with post-hoc Tukey’s multiple comparison test).

Increased dFB activity is thought to drive increased sleep in juvenile flies (Kayser et al., 2014), leading us to ask whether dFB inhibition in juvenile flies result in a mature-like sleep state. We drove expression of the inwardly-rectifying potassium channel, *Kir2*.*1*, in *R23E10-GAL4* neurons. To account for developmental effects of inhibiting the dFB, we utilized a ubiquitously-expressed temperature-sensitive *GAL80* repressor protein (*tub-GAL80*^*ts*^) (McGuire et al., 2004). Raising the temperature rapidly degrades *GAL80*^*ts*^, expressing the downstream *UAS* transgene. In juvenile flies, expressing a GFP-tagged Kir2.1 in *R23E10-GAL4* neurons (*tub-GAL80*^*ts*^; *R23E10-GAL4>UAS-Kir2*.*1*) significantly decreased sleep duration during the night (**Fig 4A-C**). Sleep/wake transition probabilities were unaffected with dFB inhibition in juvenile flies (**Fig 4D-E**); however, nighttime deep sleep was decreased, while light sleep and light wake increased (**Fig 4F**). Thus, dFB inhibition in juvenile flies did not affect fully reflect mature-like sleep architecture. Together, these results suggest the dFB regulates different aspects of sleep architecture in mature and juvenile flies.

**Fig 4:**
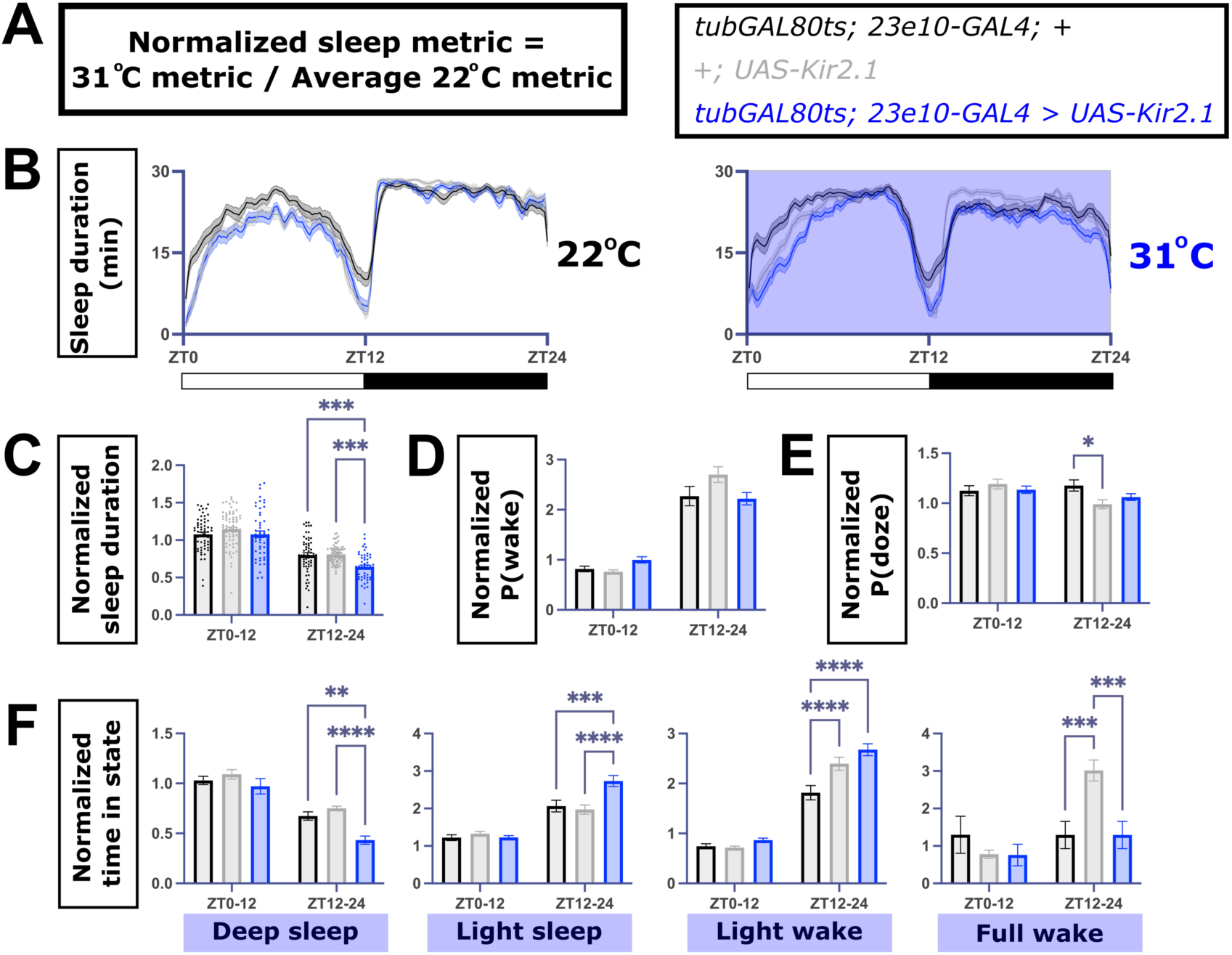
Acutely expressing *Kir2*.*1* in *R23E10-GAL4+* neurons decreases sleep duration in juvenile flies but does not recapitulate mature fly sleep architecture. A) Formula used to calculated normalized sleep metric. Sleep was recorded in juvenile flies one day post-eclosion, and sleep metrics were normalized to the average of the baseline at 22°C. Sleep duration traces of B) juvenile *tubGAL80ts; R23E10-GAL4>UAS-Kir2*.*1* (blue) vs genetic controls (black and gray) at 22°C (left) and 31°C (right). Normalized C) sleep duration, D) P(wake), E) P(doze), and F) time spent in each sleep state in juvenile flies (n = 75, 60, 54 from left to right, one-way ANOVA with post-hoc Tukey’s multiple comparison test).

### Distinct molecular profiles in juvenile vs mature dorsal fan-shaped body neurons reflects differential sleep-regulatory functions across the lifespan

Maturation of dopaminergic projections to the dFB is a key event for sleep ontogeny (Chakravarti Dilley et al., 2020; Kayser et al., 2014), but whether sleep-promoting dFB neurons undergo intrinsic maturation is unknown. Single-cell RNA-Seq analysis of the adult fly brain at different ages previously identified a cluster of cells that contain those matching the expression profile of *R23E10-GAL4* sleep-promoting neurons (Davie et al., 2018). This cluster exhibited 55 differentially expressed genes (DEGs) between mature (day 9 post-eclosion) and juvenile (day 0-1 post-eclosion) flies (**Fig S3A**) (Davie et al., 2018). We used this dataset to ask how the transcriptomic profiles of dFB cells change during development. First, to identify mechanisms that might be responsible for dFB function in juvenile and mature flies, we performed gene set enrichment analysis (GSEA; Mootha et al., 2003; Subramanian et al., 2005). GSEA revealed the DEGs that were more highly expressed in mature flies were enriched for ribosomal and translational processes (**Table S2**). Conversely, while DEGs that were more highly expressed in juvenile flies were not significantly enriched for specific processes, we noted several of these genes were involved in transmembrane ion transport, synaptic transmission, and neurodevelopment (**Table S3**). Thus, dFB cells exhibit distinct gene expression profiles in juvenile and mature flies.

**Table S2:** GSEA results for DEGs with increased expression in mature compared to juvenile dFB cells.

| Gene set name | # genes in gene set | Enrichment score | Normalized enrichment score | Nominal p-value | FDR q-val |
| --- | --- | --- | --- | --- | --- |
| RIBONUCLEOPROTEIN COMPLEX | 25 | 0.8334325 | 3.3576221 | 0 | 0 |
| STRUCTURAL CONSTITUENT OF RIBOSOME | 23 | 0.7740875 | 3.0646384 | 0 | 0 |
| RIBOSOME | 23 | 0.7740875 | 3.0391607 | 0 | 0 |
| CYTOSOLIC PART | 23 | 0.7740875 | 3.0359354 | 0 | 0 |
| STRUCTURAL MOLECULE ACTIVITY | 23 | 0.7740875 | 3.0340197 | 0 | 0 |
| RIBOSOMAL SUBUNIT | 23 | 0.7740875 | 3.0132124 | 0 | 0 |
| CYTOSOLIC RIBOSOME | 23 | 0.7740875 | 3.0076745 | 0 | 0 |
| MITOTIC CELL CYCLE PROCESS | 15 | 0.5640415 | 2.0024745 | 0.00466201 | 0.00371607 |
| MICROTUBULE CYTOSKELETON ORGANIZATION | 15 | 0.5558247 | 1.9938301 | 0.00943396 | 0.00354415 |
| MICROTUBULE BASED PROCESS | 15 | 0.5558247 | 1.9627372 | 0.00485437 | 0.00352023 |
| CYTOSKELETON ORGANIZATION | 15 | 0.5558247 | 1.9592851 | 0 | 0.00320021 |
| MITOTIC CELL CYCLE | 16 | 0.5014893 | 1.7472951 | 0.01678657 | 0.01589365 |
| PROTEIN COMPLEX SUBUNIT ORGANIZATION | 15 | 0.46264157 | 1.6513298 | 0.02849741 | 0.02640818 |

**Table S3:**
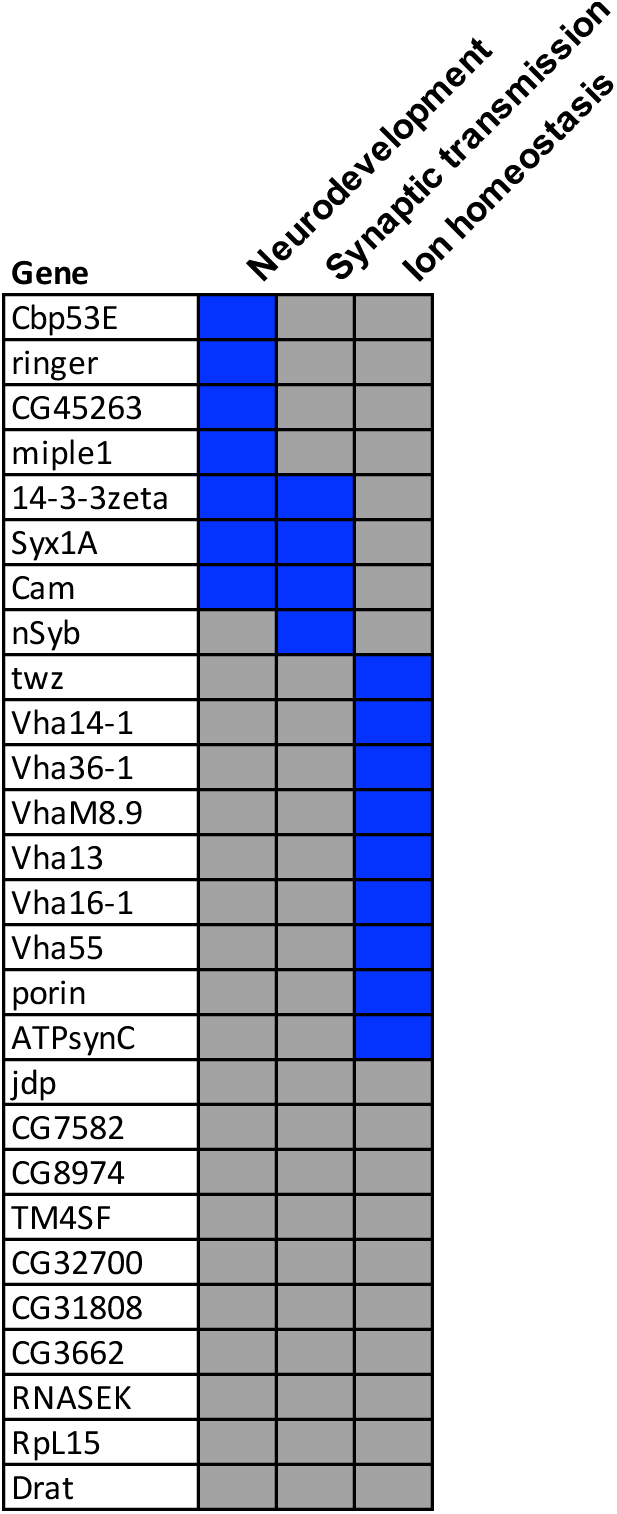
Associated GO terms for DEGs with increased expression in juvenile fly dFB cells. Blue boxes indicate a given gene is associated with the listed GO term, while gray indicates the gene is not associated with a GO term.

Next, we sought to determine whether developmental changes in the molecular landscape of *R23E10-GAL4+* neurons relate to intrinsic maturation of these cells. We reasoned that DEGs with higher expression in the juvenile compared to mature dFB cells could be involved in the maturation of this sleep center, such that knockdown of these genes would affect sleep in the mature fly but not the juvenile fly. Specifically, we hypothesized sleep-promoting dFB neurons would be stunted in a more juvenile state. Using the *R23E10-GAL4* driver, we individually knocked down genes that were more highly expressed in the juvenile dFB neurons and recorded sleep in both juvenile and mature flies. When compared to genetic controls (*R23E10-GAL4>UAS-mCherry RNAi*), knockdown of DEGs with increased expression in juvenile flies did not differentially affect sleep duration from ZT0-12 (when juvenile and mature flies exhibited the largest differences in sleep duration) in juvenile versus mature flies (**Fig 5A; Fig S3B**). However, as our work demonstrates, focusing solely on sleep duration fails to capture more nuanced differences in sleep states between juvenile and mature flies. To examine sleep states, we first trained a HMM on data from mature *R23E10-GAL4>UAS-mCherry RNAi* control flies to account for genetic background. These flies exhibited the same differences in sleep/wake transition probabilities (**Fig S4A)** and HMM substates (**Fig S4B**) as *iso31* flies, indicating these changes remain consistent across genetic background. To determine whether knockdown of juvenile DEGs affects sleep states, we focused on P(wake) (**Fig 5B; Fig S3C**) and deep sleep **(Fig 5C; Fig S3D**) at ZT0-12, as these are the metrics with the largest differences between mature and juvenile flies. Compared to age-matched genetic controls, knockdown of genes with higher expression in juvenile dFB cells was associated with increased deep sleep in mature flies versus juvenile flies (**Fig 5Ci**). This result is consistent with our hypothesis that knockdown of juvenile-specific dFB genes results in persistent immaturity of dFB function in the mature fly. Conversely, knockdown of genes with higher expression in mature dFB cells did not differentially affect sleep duration (**Fig 5Aii)**, P(wake) (**Fig 5Bii**), or deep sleep (**Fig 5Cii**) across age groups. These results provide functional evidence that distinct biological processes present in juvenile fly dFB cells are important for *R23E10-GAL4* neuron maturation.

**Fig 5:**
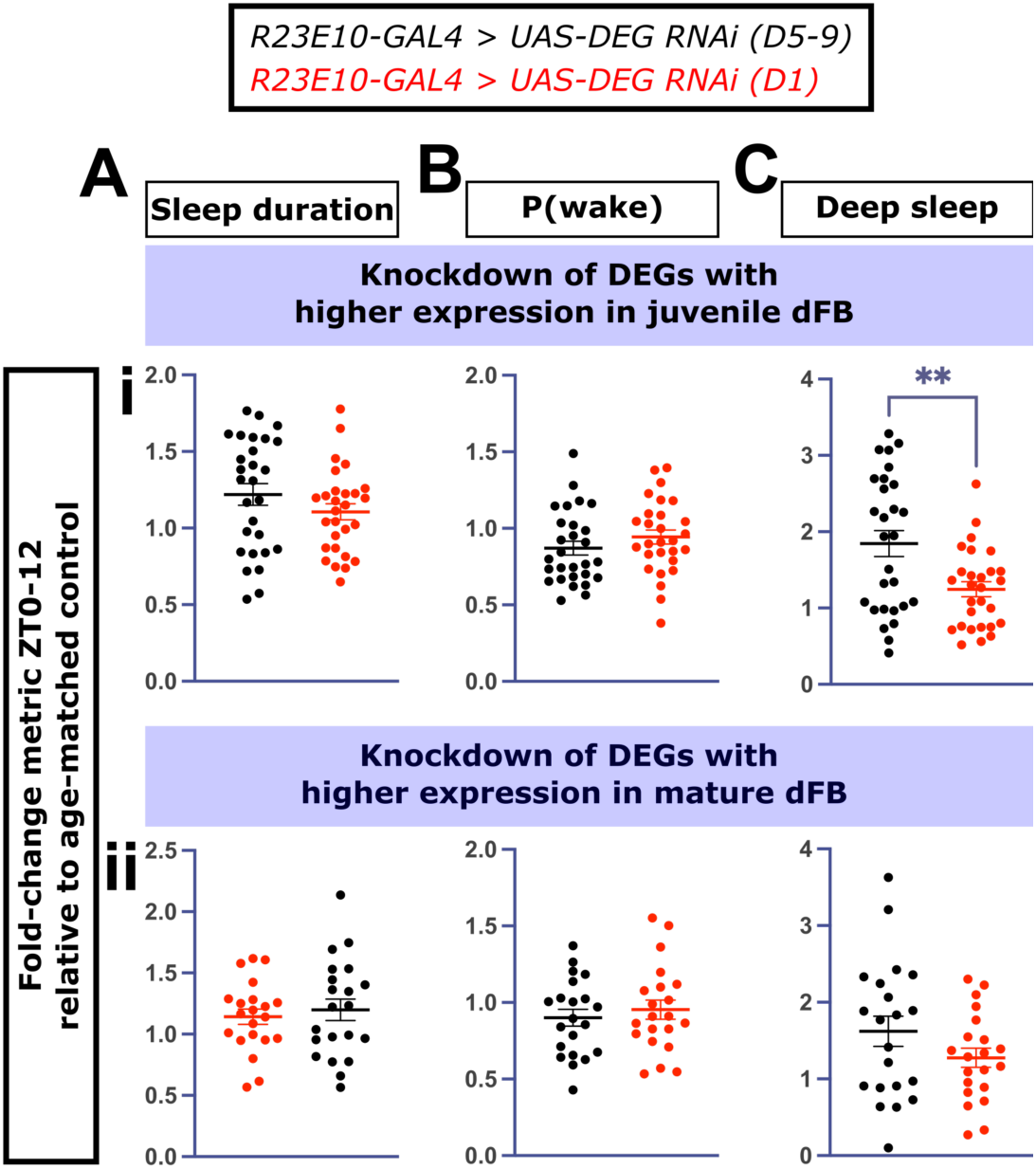
Knockdown of DEGs with higher expression in juvenile dFB cells increases deep sleep in mature flies more than in juvenile flies. Fold-change in A) sleep duration, B) P(wake) and C) deep sleep from ZT0-12 in mature (black) and juvenile (red) flies compared to age-matched genetic controls (*R23E10-GAL4>UAS-mCherry RNAi)* in the setting of *R23E10-GAL4*-mediated knockdown of overexpressed genes in i) juvenile dFB cells and ii) mature dFB cells. Each data point represents a different RNAi line for a specific DEG; n ≥ 10 flies per line (two-tailed T-tests).

**Fig S3:**
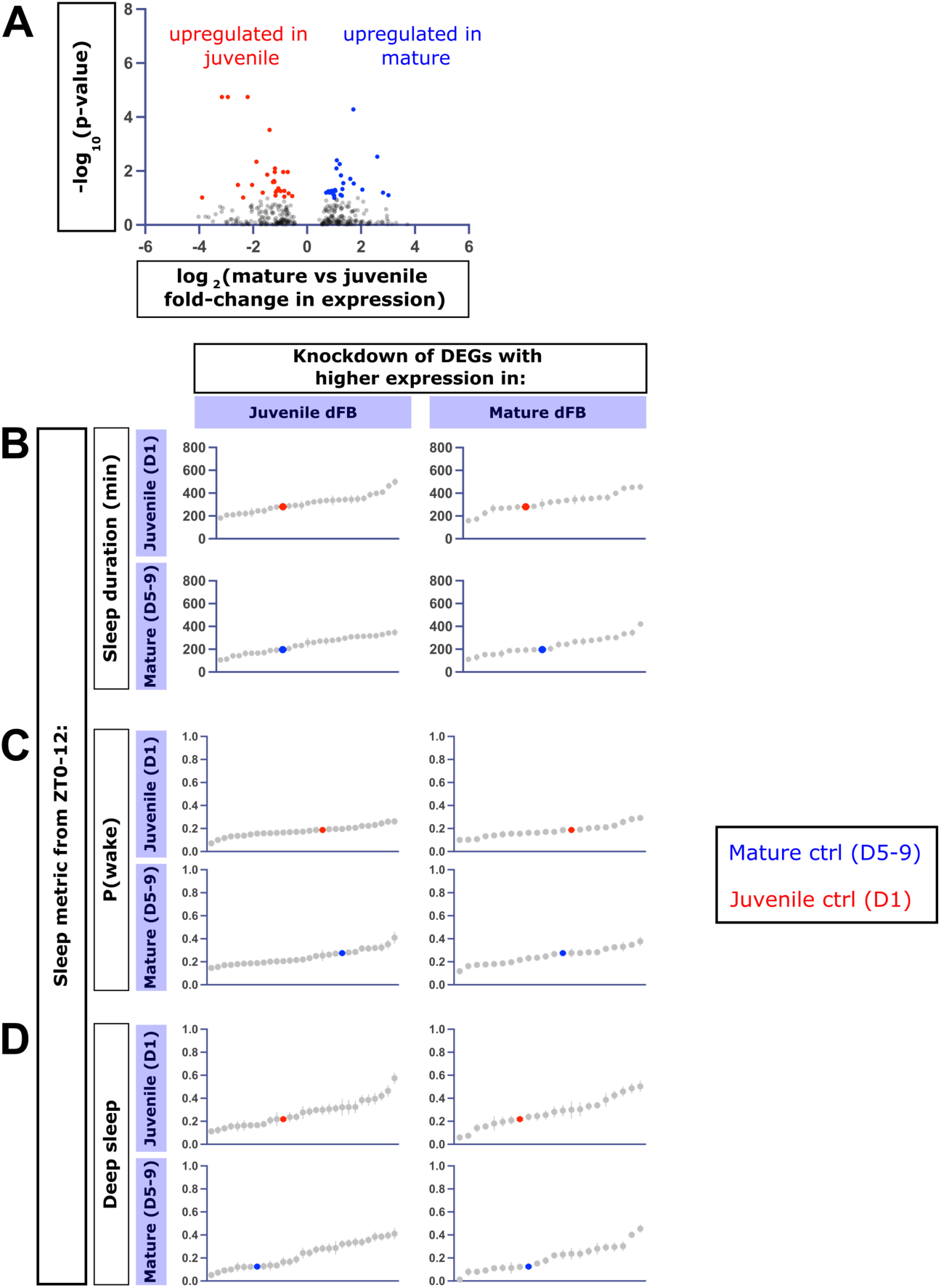
Sleep duration, P(wake), and proportion of time spent in deep sleep from an RNAi-based screen of differentially-expressed genes in dFB neurons in juvenile vs mature flies. A) Differentially expressed genes (DEGs) based on published datasets (Davie et al., 2018). Red: genes that are more highly expressed in juvenile vs mature flies, blue: genes that are more highly expressed in mature vs juvenile flies, based on an adjusted p-value cut-off (p-adj > 0.1). B) Sleep duration, C) P(wake), and D) proportion of time in deep sleep across ZT0-12 in juvenile vs mature flies in *R23E10-GAL4>UAS-RNAi* (n ≥ 10 per RNAi line; gray) compared to age-matched genetic controls (juvenile: red; mature: blue). Knockdown of DEGs with higher expression in juvenile dFB neurons (left graphs in B,C) result in comparable sleep duration and P(wake) distributions in juvenile vs mature flies around the age-matched control. Knockdown of genes with higher expression in juvenile dFB neurons skews mature fly deep sleep to the right (D, bottom left) compared to knockdown of the same genes in juvenile flies (D, top left). Knockdown of DEGs with higher expression in mature dFB neurons (right graphs in B-D) does not differentially skew sleep metric distributions when comparing juvenile vs mature flies.

**Fig S4:**
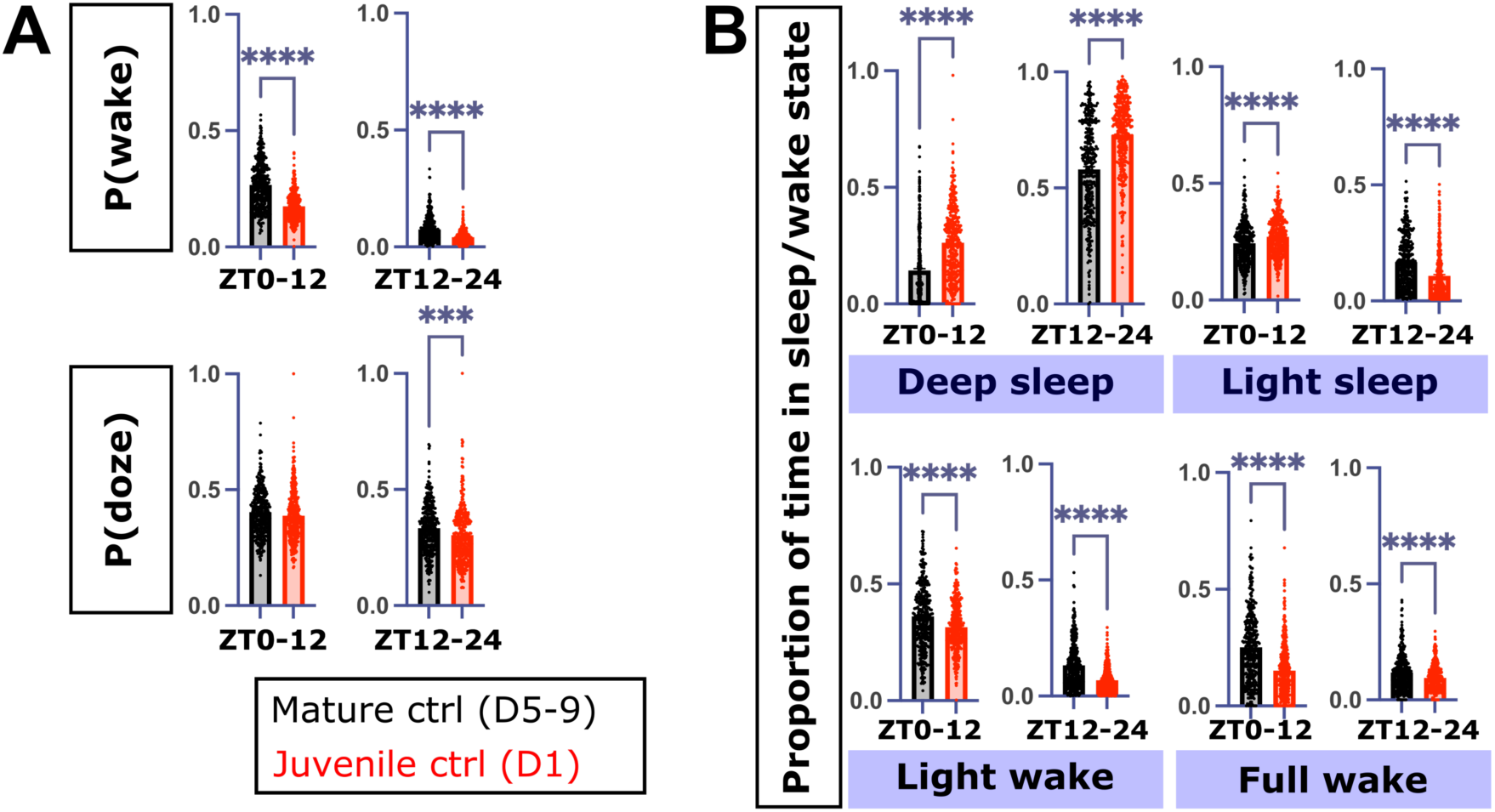
*R23E10-GAL4>UAS-mCherry RNAi* control flies exhibit the same sleep architecture differences across development as *iso31* flies. A) P(wake) (top), P(doze) (bottom), and B) proportion of time spent in sleep/wake states from ZT0-12 and ZT12-24 in mature (black, n = 344) vs juvenile (red, n = 347) *R23E10-GAL4>UAS-mCherry RNAi* controls (two-tailed T-tests).

## Discussion

Sleep duration in early life is consistently greater than later in life across species, but how maturation of individual neural circuits contributes to ontogenetic changes in sleep architecture is unclear. In this study, we describe changes in sleep/wake transition probabilities and substates in *Drosophila* that accompany changes in sleep duration across the lifespan. Using these probabilistic methods, we identify mechanisms underlying intrinsic dFB maturation contributing to sleep maturation. Our results link changes in the molecular profile of sleep output neurons to sleep ontogeny.

Here, we demonstrated quantifiable differences in sleep architecture across the lifespan. How does the unique sleep quality in juvenile flies contribute to neurodevelopment? In mammals, REM and non-REM sleep differentially contribute to development (Knoop et al., 2021). The proportion of REM sleep is significantly increased in neonates (Roffwarg et al., 1966), and plays a critical role in neurodevelopment. For example, REM sleep is necessary for plasticity in the developing visual cortex (Bridi et al., 2015; Frank et al., 2001; Shaffery et al., 2002) and selective strengthening of synaptic contacts occurs during REM sleep in early life (Li et al., 2017). A preponderance of motor twitches also occurs during REM sleep in young animals, and increased REM is thought to be important for patterning of sensorimotor circuits from these inputs (Blumberg et al., 2013; Mohns and Blumberg, 2010; Sokoloff et al., 2021). Despite evidence that REM sleep is important for neurodevelopment, we understand little about the genetic mechanisms linking the two. Furthermore, non-REM sleep is proportionally decreased compared to REM sleep, but still plays a role in synaptic pruning (Tononi and Cirelli, 2006) and cortical maturation (Kurth et al., 2010), especially during later developmental periods beyond the neonatal stage. However, like with REM sleep, the mechanisms connecting NREM sleep to neurodevelopment remain unknown. Our study establishes a genetically-tractable model to identify molecular regulators of sleep states that are important for sleep-dependent neurodevelopment.

While development of arousal-promoting dopaminergic neurons is known to be essential for normal sleep ontogeny (Chakravarti Dilley et al., 2020), we now find that intrinsic maturation in sleep output neurons also contributes to differences in sleep between mature and juvenile animals. The dFB exhibits increased activity in juvenile compared to mature flies, which results in excess sleep duration in early life (Kayser et al., 2014). We found while dFB inhibition decreases daily sleep in juvenile flies, this manipulation does not result in “mature-like” sleep architecture. Conversely, even though dFB activation in mature flies increases sleep duration, this sleep does not fully recapitulate the juvenile sleep state from a sleep architecture perspective. In mature flies, the dFB is involved in rebound following sleep deprivation, and disrupting the function of the dFB by knocking down various signaling components blunts rebound (Donlea et al., 2018, 2014; Qian et al., 2017). These results suggest that the dFB regulates sleep during periods of increased homeostatic drive, such as during early life and in the sleep-deprived mature adult. However, several lines of evidence suggest sleep in juvenile flies is nonetheless distinct compared to rebound sleep in mature flies (Dilley et al., 2018). Of note, a recent study showed that sleep resulting from dFB activation in mature flies is electrophysiologically-distinct from endogenous rebound sleep in mature flies (Tainton-Heap et al., 2021). We additionally show rebound sleep in mature flies is distinct from juvenile fly sleep, indicating juvenile fly sleep is not simply the same state of heightened homeostatic drive. Several circuits may also act together to differentially influence sleep architecture in juvenile versus mature flies, which may explain the differences we see with dFB manipulation. Nevertheless, single-cell RNA Seq analysis revealed distinct molecular profiles in the dFB in juvenile flies compared to mature flies, supporting the hypothesis that these neurons undergo intrinsic development that may govern differential sleep-regulatory function in juvenile and mature flies. Our functional studies suggest the sleep-promoting dFB neurons have a changing role in sleep across development: while they influence baseline sleep in juvenile flies, they play a more specific role in rebound sleep in mature flies.

We show that distinct molecular mechanisms present in juvenile fly dFB cells govern dFB function maturation, but how these processes are involved in the development of the dFB in the context of sleep ontogeny is unclear. Knockdown of ribosomal function and translation-related DEGs that were overexpressed in the mature fly dFB neurons did not differentially affect sleep in juvenile versus mature flies. Conversely, knockdown of DEGs involved in synaptic function, ion homeostasis, and neurodevelopment that were overexpressed in juvenile fly dFB increased deep sleep in the mature fly more so than in juvenile flies. Notably, knockdown of these DEGs did not differentially affect sleep duration in mature and juvenile flies, even though we observed significant effects on sleep architecture. These results highlight the utility of non-invasive computational approaches in the fly for investigating sleep architecture. One possible interpretation of these findings is that genes with higher expression in dFB in early adult life are important for the maturation of *R23E10-GAL4+* neurons, while genes that are more highly expressed later in life are important for the sleep-regulatory function of these neurons in mature flies. For example, genes with higher expression in the mature dFB neurons may be involved in mediating appropriate sleep rebound following deprivation. Another possibility lies in the heterogeneity of dFB sleep neurons: individual dFB neurons exhibit vastly different excitabilities (Donlea et al., 2014; Pimentel et al., 2016), suggesting the dFB contains a diverse group of sleep neurons.

Knockdown of genes that are overexpressed in juvenile flies may inhibit the development of dFB neurons that are specifically relevant in mature flies. Intersectional approaches to investigate the contribution of specific sub groups of dFB neurons to sleep in juvenile and mature flies would be informative for our understanding of the dFB circuits underlying sleep ontogeny. Together, these results provide a framework for understanding the molecular processes governing maturation of sleep output neurons to influence sleep ontogeny.

## Materials and Methods

### Fly stocks

Flies were raised and maintained on standard molasses food (8.0% molasses, 0.55% agar, 0.2% Tegosept, 0.5% propionic acid) at 25 °C on a 12hr:12hr light:dark cycle. Female flies were used in all experiments.

### Fly strains

*Iso31* was a laboratory strain. *UAS-dTrpA1* was a gift from Dr. Leslie Griffith (Brandeis University). *R23E10-GAL4, UAS-Kir2*.*1-GFP*, and *UAS-mCherry RNAi* were obtained from the Bloomington Drosophila Resource Center. All RNAi strains were obtained from Bloomington Drosophila Resource Center.

### Sleep assays

For ontogeny experiments unless otherwise specified, newly-eclosed female flies were collected and aged in group housing on standard food. Juvenile flies were collected on the day of eclosion and loaded into the DAM system between ZT4-6, along with mature flies aged 5-9 days post-eclosion. Unless otherwise specified, sleep assays were run at 25 °C on a 12-hour/12-hour light/dark schedule.

### Thermogenetic activation and inhibition experiments

Animals were raised at 18 °C to prevent TrpA1 activation or Kir2.1 expression during development. For TrpA1 activation experiments, adult female flies were collected 2-3 days post-eclosion and aged at 18 °C on standard fly food. 5-9 day old flies were loaded into the DAM system to monitor sleep and placed at 22 °C on a 12:12: LD schedule for 3 days. TrpA1 activation was performed by a temperature shift to 31 °C across non-consecutive 12-hour light or 12-hour dark periods. Between increases in temperature, flies were maintained at 22 °C. For Kir2.1 inhibition experiments, adult female flies were collected at eclosion and aged at 18 °C in group-housed conditions. Juvenile flies were collected at eclosion from ZT4-6 and loaded into the DAM system along with 5-9 day old flies at 31 °C. For Kir2.1-GFP immunohistochemistry experiments, flies were collected as described above and shifted to 31 °C 20 hours before dissection.

### Sleep/wake transition probabilities and hidden Markov modeling analysis

P(wake) and P(doze) were calculated from 1-minute bins of activity collected in the DAM system in Matlab as previously described (Wiggin et al., 2020). Hidden Markov modeling of sleep/wake substates was constrained with parameters as previously described (Wiggin et al., 2020): a transition from deep sleep to full wake could only do so through light wake, while a transition from full wake to deep sleep could only do so through light sleep. HMMs were trained on the transitions (wake or doze) between 1-minute bins of activity (for 24 hours, 1439 transitions per fly). HMM fitting and hidden state analysis was performed as previously described using the Matlab Statistics and Machine Learning Toolkit (Wiggin et al., 2020). For characterizing ontogenetic differences in juvenile vs mature *iso31* fly sleep/wake substates (**Fig 1** and associated supplemental figures), HMMs were trained based on transitions as measured using the DAM5H multibeam system (Trikinetics). For *iso31* sleep deprivation experiments (**Fig 2**) and *R23E10-GAL4+* neuron functional manipulations (**Fig 3-4**), an HMM was trained on mature *iso31* activity transitions measured using the single beam DAM system (Trikinetics) (see **Table S4** for transition and emission probabilities). A separate HMM was trained on activity transitions measured using the single beam DAM system from all *R23E10-GAL4>UAS-mCherry RNAi* flies (see **Table S5** for transition and emission probabilities). Trained HMMs were used to calculate the proportion of time spent in sleep/wake hidden states.

**Table S4:**
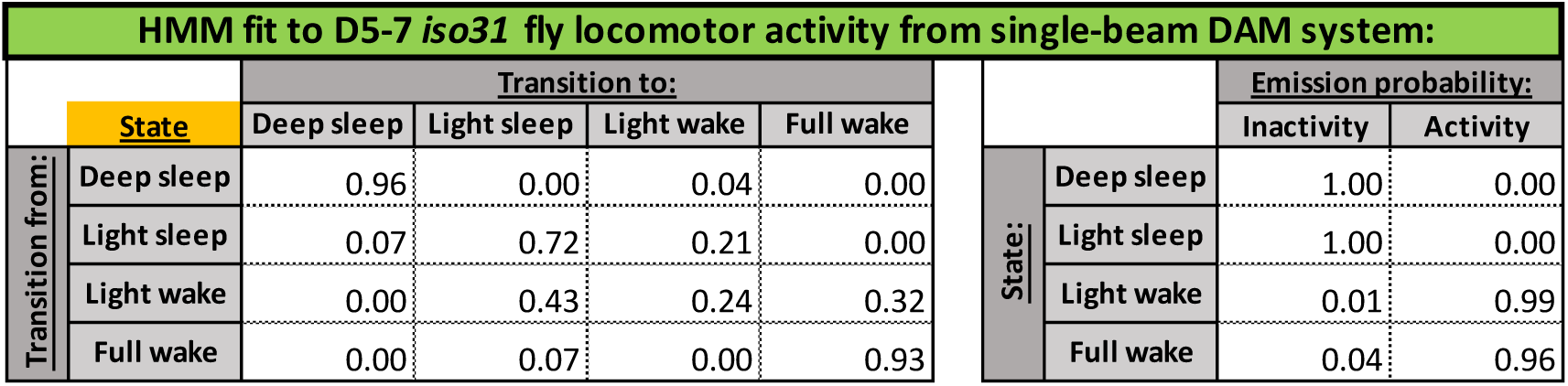
HMM parameters used to calculate proportion of time spent in sleep/wake substates for Figures 2-4. Transition probabilities between hidden states and emission probabilities from each hidden state to observed states for HMM trained on mature *iso31* fly (n = 90 flies) locomotor data collected using the single beam DAM system.

**Table S5:**
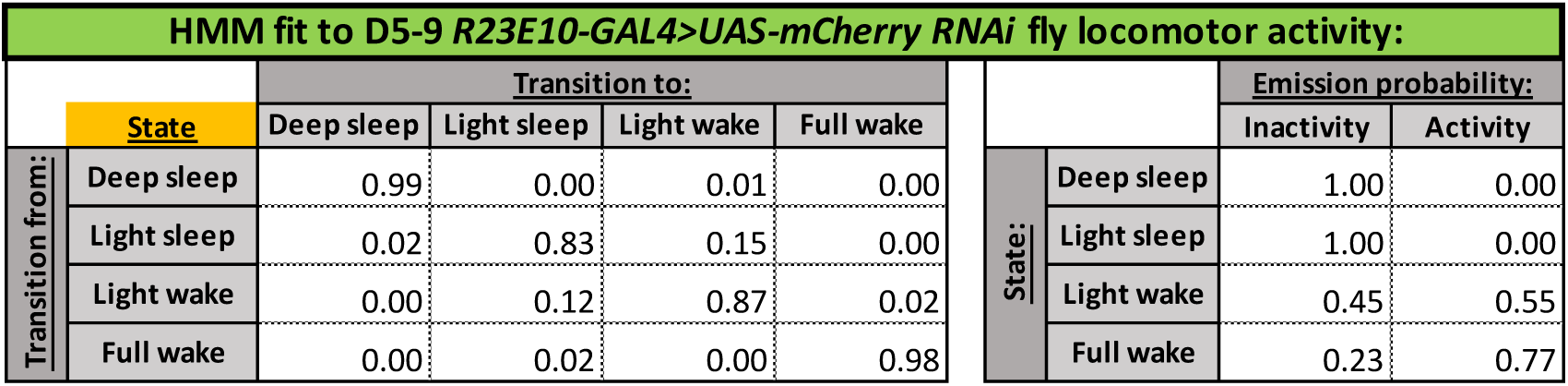
HMM parameters used to calculate proportion of time spent in sleep/wake substates for Figure 5 and associated supplemental figures. Transition probabilities between hidden states and emission probabilities from each hidden state to observed states for HMM trained on mature *R23E10-GAL4>UAS-mCherry RNAi* fly (n = 344 flies) locomotor data collected using the single beam DAM system

#### Single cell RNA-Seq analysis

Using published single-cell data from *Drosophila* brains, cells annotated as dFB neurons were extracted from previously performed clustering(Davie et al., 2018) (cluster 61 in the 57K dataset with clustering resolution 2.0) and collapsed into pseudobulk transcriptomes per replicate. Differential expression comparing young (d0 or d1 flies) vs old (d9 flies) was performed on both sets of pseudobulk transcriptomes using DESeq2 (Love et al., 2014). Genes significantly differentially expressed (p-adj < 0.1) formed the candidate list for the RNAi-based screen.

### Gene set enrichment analysis of differentially expressed dFB genes

Gene set collections for Gene Ontology annotations were downloaded from public sources (Powell, 2014). To compare DEGs upregulated in mature or juvenile flies, a gene signature was generated by ranking all DEGs with p-adj > 0.1 according to DEseq2-derived test statistics. Enrichment analysis was performed with GSEA v4.0 (Subramanian et al., 2005) using weighted statistical analysis. Gene sets with a false discovery rate < 0.25 were considered significantly enriched.

### RNAi-based ontogeny screen of differentially expressed dFB genes

Virgin collected from the *R23E10-GAL4* fly stock were crossed to males of RNAi fly stocks from the Transgenic RNAi Project (TRiP) collection (Ni et al., 2011). We utilized all available VALIUM10, VALIUM20, or VALIUM22 lines for a given gene. For controls, we used *R23E10-GAL4* x *UAS-mCherry RNAi*. Sleep ontogeny assays were performed as described above. The DAM system was used to collect 1-minute bins of activity for calculating sleep/wake transition probabilities and HMM hidden states.

### Statistical analysis

All statistical analyses were performed using GraphPad Prism (version 8.4.1). Sample size, specific tests, and significance values are denoted in figure legends.

## Acknowledgements

We thank members of the Kayser Lab, Raizen Lab, Leela Chakravarti Dilley, and members of the Penn Chronobiology and Sleep Institute for helpful discussions/input. We thank Leslie Griffith and Timothy Wiggin for assisting with the custom Matlab codes used for sleep/wake transition probabilities and hidden Markov modeling of sleep/wake substates.

